# Lipid metabolic reprogramming drives triglyceride storage and variable sensitivity to FASN inhibition in endocrine-resistant breast cancer cells

**DOI:** 10.1101/2024.05.30.596684

**Authors:** Ashley V. Ward, Duncan Riley, Kirsten E. Cosper, Jessica Finlay-Schultz, Heather M. Brechbuhl, Andrew E. Libby, Kaitlyn B. Hill, Rohan R. Varshney, Peter Kabos, Michael C. Rudolph, Carol A. Sartorius

## Abstract

**Background:** Lipid metabolic reprogramming is increasingly recognized as a hallmark of endocrine resistance in estrogen receptor-positive (ER+) breast cancer. In this study, we investigated alterations in lipid metabolism in ER+ breast cancer cell lines with acquired resistance to common endocrine therapies and evaluated the efficacy of a clinically relevant fatty acid synthase (FASN) inhibitor.

**Methods:** ER+ breast cancer cell lines resistant to Tamoxifen (TamR), Fulvestrant (FulvR), and long-term estrogen withdrawal (EWD) were derived. Global gene expression and lipidomic profiling were performed to compare parental and endocrine resistant cells. Lipid storage was assessed using Oil Red O (ORO) staining. The FASN inhibitor TVB-2640 was tested for its impact on lipid storage and cell growth. ^13^C_2_-acetate tracing was used to evaluate FASN activity and the efficacy of TVB-2640.

**Results:** Endocrine resistant cells showed significant enrichment in lipid metabolism pathways and distinct lipidomic profiles, characterized by elevated triglyceride levels and enhanced cytoplasmic lipid droplets. ^13^C_2_-acetate tracing revealed increased FASN activity in endocrine resistant cells, which was effectively reduced by TVB-2640. While TVB-2640 reduced lipid storage in most but not all cell lines, this did not correlate with decreased cell growth.

Polyunsaturated fatty acids (PUFAs) containing 6 or more double bonds were elevated in endocrine resistant cells and remained unaffected or increased with TVB-2640.

**Conclusion:** Endocrine resistant breast cancer cells undergo a metabolic shift toward increased triglyceride storage and PUFAs with high degrees of desaturation. While TVB-2640 reduced lipid storage in most conditions, it had limited effects on the growth of endocrine resistant breast cancer cells. Targeting specific lipid metabolic dependencies, particularly pathways that produce PUFAs, represents a potential therapeutic strategy in endocrine resistant breast cancer.

## Background

Overcoming resistance to endocrine therapies represents a pivotal challenge for enhancing treatment outcomes in women with estrogen receptor-alpha (ER) positive breast cancer, contributing to approximately half of all fatalities from the disease (1). The three most common classes of drugs used during endocrine therapy include selective ER modulators (SERMS, e.g., Tamoxifen, (Tam)) that disrupt binding of estrogens to ER, selective ER degraders (SERDs, e.g., Fulvestrant (Fulv)) that cause immobilization and/or degradation of ER (2), and aromatase inhibitors that block production of 17β-estradiol. Typically administered as adjuvant therapy for over 5-10 years, endocrine therapy offers long-term disease stabilization or eradication for most patients. Unfortunately, acquired endocrine resistance emerges in up to one third of women either during treatment or upon its conclusion, often coinciding with distal metastasis (3). At this time, several targeted therapies may be integrated in combination with endocrine therapy, including cyclin dependent kinases 4/6 inhibitors (i.e. palbociclib), alpelisib for PIK3CA mutations, or the SERD elacestrant for tumors harboring ER mutations (4, 5).

However, the development of acquired multidrug resistance is usually inevitable. Consequently, an urgent need exists to identify new vulnerabilities in endocrine resistant cells that can be effectively targeted in combination with endocrine drugs to mitigate the recurrence risk or delay progression of recurrent disease.

ER+ breast cancers exhibit pronounced lipogenesis, utilizing de novo fatty acid (FA) synthesis despite the presence of abundant FAs in their microenvironment (6). FAs originating from preformed sources facilitate synthesis of more complex lipids necessary for membrane expansion, provide fuel for rapid tumor growth, and act as signaling molecules. Central to this process is the upregulation of Sterol Regulatory Element-binding transcription factor 1 (SREBP1) by estrogens, which in turn boosts the expression of two vital de novo FA synthesis enzymes necessary in this pathway: acetyl-CoA carboxylase (ACC) and fatty-acid synthase (FASN) (7). ACC catalyzes the conversion of acetyl-CoA into malonyl-CoA, and both precursors are needed for further catalysis by FASN to synthesize palmitate. Palmitate and further elongated FAs are subsequently combined with preformed FA taken up from the microenvironment by transporters, and together, they are subsequently incorporated into triglycerides and frequently stored as cytoplasmic lipid droplets (LD) in breast cancer cells (8–10). This dual mode of FA synthesis and FA uptake underscores the complexity of targeting FA dependence in breast cancers, especially given the adaptability in the metabolic incorporation of these essential molecules that provide an advantage to cancer cells.

Recent studies have shown that lipid metabolic pathways undergo further alteration with the development of endocrine resistance, with a significant body of research centering on Tam, the longest approved breast cancer drug (11). Hultsch et al. observed that Tam-resistant breast cancer cells had a significant increase in triglycerides and cholesterol esters, stored in large LDs and lysosomes, respectively (12). Moreover, ER+ invasive lobular carcinoma cells, resistant to long term estrogen deprivation, exhibited heightened activation of lipid metabolism that could be stratified into two distinct lipid pathways, SREPB1-activated FA synthesis or cholesterol metabolism (13). Notably, MCF7 cells subjected to long-term estrogen deprivation experience an uptick in cholesterol biosynthesis (14). HER2-overexpressing MCF7 variants had elevated FASN expression alongside an increased dependency on lipid biosynthesis (7).

While multiple potential targets within lipid metabolism are promising, FASN inhibition has historically received the most attention. FASN levels are consistently higher in breast cancer compared to normal cells and breast tumor vs normal tissue (15). While early-generation FASN inhibitors demonstrated potential efficacy for tumor growth restriction in preclinical breast cancer models, their clinical translation was limited due to substantial toxicity observed in both animal models and human trials (16). However, a more recently developed selective and reversible FASN inhibitor (TVB-2640) has shown promising results, including dose-dependent efficacy in advanced solid tumors like HER2-positive breast cancers and has advanced to Phase II clinical trials (17). Beyond FASN, other targets within lipid metabolism are currently being explored, such as inhibitors of fatty acid oxidation (FAO), fatty acid transport, and the synthesis of monounsaturated fatty acids (18). Given the possibility for compensation within lipid metabolic pathways, therapeutic strategies targeting single enzymes are usually met with metabolic adaptations, suggesting combinatorial approaches using established drugs might yield stronger outcomes.

In this study, we analyzed the global lipidome of breast cancer cells exhibiting resistance to three common endocrine therapy modalities. Our findings reveal that while the lipid composition of endocrine resistant cells is distinct from each other and from their endocrine sensitive counterparts, they share an increased abundance of triglycerides, LDs and complex lipids containing highly polyunsaturated FA (PUFA) species. The FASN inhibitor TVB-2640 effectively blocked FASN activity, but showed variable effects on lipid storage and cell growth, with only modest reductions observed in some resistant subtypes. Furthermore, certain endocrine treatments enhanced PUFA levels when FASN was inhibited. These results underscore the distinct lipid metabolism signature of endocrine resistant breast cancer cells and highlight the challenges of targeting FASN in endocrine resistance. Importantly, a reliance on lipid desaturation pathways emerges as a potential vulnerability in endocrine resistance.

## Materials and Methods

### Generation of Endocrine Therapy Resistant Cell Lines

T47D cells were maintained in minimal Eagle’s medium supplemented with 5% fetal bovine serum (FBS), 1x non-essential amino acids (NEAA), 1 nM insulin, and 1 mM sodium pyruvate unless otherwise noted. The generation and characterization of breast cancer patient-derived xenograft (PDX)- derived cell lines UCD4 and UCD65 was previously described (19). UCD4 and UCD65 cells were maintained in Dulbecco’s Modified Eagle Media/F-12 (50:50) supplemented with 10% FBS, 100 ng/mL choleratoxin, and 1 nM insulin. Cell lines were authenticated by short tandem repeat analysis and routinely tested for mycoplasma using the MycoAlert mycoplasma detection kit (Lonza, #LT07-710). UCD4 contains an ESR1 mutation (D538G).

To establish endocrine resistant variants, cells were exposed to escalating doses of ICI 182780 (Fulv, Faslodex, T47D and UCD4) or 4-hydroxytamoxifen (4OHT, Tam, T47D, UCD4, and UCD65) over a 12-month period to establish endocrine resistance. The starting dose for Fulv and Tam were 10 times lower than the effective dose for each compound, 1 nM and 10 nM respectively. Both compounds were dissolved in 100% EtOH to make stock concentrations that resulted in the final dose being less than 0.1% EtOH. Fulv resistant (FulvR) and Tam resistant (TamR) cells were maintained at established resistance concentrations, 100 nM and 1 µM, respectively. For estrogen withdrawal, UCD4 cells (UCD4-EWD), were cultured for 12-months in phenol-red free media supplemented with charcoal stripped serum.

#### RNA-sequencing

Parental and resistant cells were plated in 6-well plates at 5x10^5^ (T47D) or 1x10^6^ (UCD4) cells/well in normal media conditions with appropriate maintenance Fulv or 4OHT drug concentrations. All samples were grown to 80% confluency then harvested until all samples were ready for processing. RNA was isolated using QIAzol lysis reagent (Qiagen, #79306) and RNAeasy Mini Kit (Qiagen, #74104) and treated with RNase-free DNase (Qiagen, #79254).

Libraries were prepared by the University of Colorado Genomics Shared Resource using Illumina TruSeq Stranded mRNA Library Prep kit (Illumina, #20020595) and sequenced using paired end 50 cycles 2 x 50 on the NovaSEQ 6000 with >30 million reads per sample obtained. Paired-end 150 nucleotide reads were aligned to the human genome version GRCh38.p13 using STAR. Downstream expression analysis was performed using Cufflinks. The resulting aligned files (BAM) were then used to infer the differentially expressed gene profiles using DESeq2.

Over representation analysis (ORA) was performed using the Gene Ontology (GO) Cellular Component gene set collection via WebGestalt (20).

### Unbiased Lipid Mass Spectrometry

1x10^6^ cells were plated in 6 cm plates in normal media conditions (N=4 per condition). Maintenance Fulv and 4OHT drug concentrations were added to resistant cell line plates. Cells were harvested at 80% confluency. Modified MTBE liquid-liquid extraction was used with a protein crash. The lipid fraction was reconstituted in 100 µL Methanol. The aqueous fraction was reconstituted in 50 µL 5% Acetonitrile in water. The lipid fraction was run in positive ionization mode on the Agilent 6560 IMMS-QTOF, in QTOF-only mode (Analytical column: Agilent Zorbax Rapid Resolution HD (RRHD) SB-C18, 1.8 µm, 2.1x100 mm; column temp 60C; Gradient elution: 0–0.5 minutes 30–70% B, 0.5–7.42 minutes 70–100% B, 7.42–9.9 minutes 100% B, 9.9–10.0 minutes 100–30% B, 10–14.6 minutes 30% B; Flow rate 0.6mL/min; Mobile phase A: water with 0.1% formic acid; Mobile phase B: 60:36:4 isopropyl alcohol:acetonitrile:water with 0.1% formic acid). Spectral peaks were extracted using Mass Hunter Profinder version B.08.00 using a Recursive Workflow and area-to-height conversion. Data were normalized using Total Useful Signal (TUS) where an abundance ratio is calculated for each compound and the average of the ratio for all compounds becomes the normalization factor. Final data are thus "normalization target abundance/run abundance” (21). The final spectral data were imported into Mass Profiler Professional (MPP) version 14.9 filtered to be present in at least 2 out of the 16 samples. Annotation was performed using our in-house Mass and Retention Time (MSRT) database, HMDB databases, Lipid Maps database, KEGG databases, and a Metiln database. MetaboAnalyst and LION were used to generate significant pathways, based on an ANOVA-computed annotated list of compounds. Only pathways with >2 compounds in the set were reported. Unsaturated lipid species were calculated from results using total abundance.

### Targeted Lipid Mass Spectrometry

For ^13^C_2_-acetate tracer studies, 4x10^5^ cells were plated in 6 well plates in normal media conditions (N=3 per condition). Maintenance Fulv and 4OHT drug concentrations were added to resistant cell line plates. Cells were treated with 10 µM TVB-2640 or DMSO for 72 hr. After 24 hr of FASN inhibition or vehicle treatment, all wells were supplemented with 250 µM 13C-2 acetate for a 48 hr tracer incubation. Cells were harvested with trypsin, pelleted, and flash frozen in liquid nitrogen. Flash frozen cell pellets were lysed in 1ml 66% methanol (vol/vol) in 0.1M potassium phosphate buffer, pH 6.8 by vigorously vortexing twice with 20 second pulses.

Reserving 10ul from these lysates, total cellular protein estimation was done using BCA assay (Thermo Fisher, #23225). The remainder of the lysates were then transferred to 13x100 mm borosilicate glass culture tubes (Fisher Scientific, #14-961-27). Lysates were acidified with 10ul of 1N HCl (prewashed with Hexanes) and vortexed briefly. Total lipids were extracted twice by adding 1mL of isooctane:ethyl acetate 3:1 (vol/vol) and once with 1mL Hexanes; for each extraction, samples were vortexed vigorously for 10-15 sec and centrifuged at 2000g for 1 minute to complete phase separation. The upper organic layer from each extraction was combined into a new 13x100 mm borosilicate glass tube. Extracted lipids were brought to dryness and resuspend in 300 ul 2,2,4-Trimethylpentane (isooctane). For the non-esterified FA (NEFA) fraction, 100 ul of the resuspended total lipids was transferred to a new borosilicate glass tube, mixed with (30 ng) blended stable isotope internal standard, taken to dryness under gaseous N_2_, and resuspended in 25ul of 1% pentafluorobenzyl bromide in acetonitrile (vol/vol), to which 25 ul of 1% diisopropylethylamine in acetonitrile (vol/vol) was added, and samples were incubated at room temperature for 30 minutes. Pentafluorbenzyl-FA derivatives were taken to dryness under gaseous N_2_ and resuspended in 120ul hexanes of injection into the GC/MS. For the total fatty acid fraction (TFA), 20ul of the original 300 uL total lipid extract was transferred to a separate Teflon lined screw cap glass tube, mixed with (66.7 ng) blended stable isotope internal standard, and taken to dryness under gaseous N_2_. TFA samples were resuspended in 500 ul of 100% ethanol to which 500 ul of 1M NaOH was added to saponify the TFA fraction at 90C for 30 minutes, followed by acidification using 550ul of 1M HCl. Saponified samples were then extracted twice with 1.5ml of Hexanes, taken to dryness under gaseous N_2_, and derivatized as above. Derivatized TFA samples were resuspended in 267ul hexanes for injection into the GC/MS. For ^13^C_2_-acetate tracer samples, 50 ul of the resuspended total lipids was transferred to a new Teflon lined screw cap glass tube, mixed with (66.7 ng) ^2^D_31_-palmitate internal standard and taken to dryness under gaseous N_2_. Samples were saponified, extracted, and derivatized as above. Derivatized samples were resuspended in 267ul hexanes for injection. For the NEFA, TFA and tracer fractions, 1ul of pentafluorbenzyl-FA derivatives was injected and data were collected on the GC-MS (8890 GC, 5977B MSD, Agilent) DB-1MS UI column (122-0112UI, Agilent) with the following run program: 80C hold for 3 minutes, 30C/minute ramp to 125C, no hold, 25C ramp to 320C and hold for 2 minutes. The flow for the methane carrier gas was set at 1.5mL/minute. Data were acquired in full scan negative chemical ionization mode to identify fatty acids of acyl chain length from 8 to 22 carbons. Peak areas of the analyte or of the standard were measured, and the ratio of the area from the analyte-derived ion to that from the internal standard was calculated (22). ^13^C_2_-acetate tracer incorporation into palmitic acid was analyzed in two carbon increments from M+2 (257 m/z) to M+16 (271 m/z) relative to the quantitative ^2^D_31_- palmitate internal standard (23–25).

### Oil Red O Staining

500 mg of Oil Red O (ORO) powder was dissolved in 10 mL of 100% propylene glycol (PEG). While heating ORO/PEG solution in 95C bead bath 90 mL of 100% PEG was added to solution in 10 mL increments with vigorous swirling in between additions. Once ORO/PEG solution cooled to room temperature and before each use, solution was filtered using a Whatman #4 filter paper. For 2D culture: 18 mm glass coverslips were sterilized in 100% ethanol overnight and placed in 6-well plates. Parental and endocrine resistant T47D or UCD4 cells were plated on glass coverslips at 5x10^4^ cells/well in normal media conditions with appropriate maintenance Fulv or 4-OH-Tam drug concentrations. Cells were cultured for 5 days at 37C and 5% CO_2_ without refreshing medium. After 5 days, cells on coverslips were washed three times with PBS and fixed in 10% buffered formalin for 10 minutes. Formalin was aspirated followed by three more washes of PBS. 1 mL of 100% PEG was added per well for 2 minutes then aspirated to waste. 1 mL of ORO/PEG solution was added to each well and incubated for 15 minutes.

ORO/PEG solution was removed to waste, and wells were washed with 60% PEG for 2 minutes. Wells were rinsed with DI H_2_O three times and counterstained with 1mL diluted 1:50 Harris Hematoxylin (Fisher Scientific, #SH30-4D) for 10 minutes. Hematoxylin was discarded and wells were washed with DI H_2_O until clear. Cells on coverslips were mounted on Fisherbrand™ ColorFrost™ Plus Microscope Slides (Fisher Scientific, #12-550-17) using tweezers and aqueous mounting medium (Electron Microscopy Sciences, #17985-10). All slides were imaged at 20X magnification (Olympus, BX40). Neutral lipid stain was quantified on ImageJ as percent area normalized to cell number (4 fields per condition).

### Immunoblots

All cell lysates for immunoblotting were harvested in RIPA buffer. 100 µg total protein per sample were separated on 7.5% SDS-PAGE gels. Primary antibodies were as follows: FASN (rabbit #3180, Cell Signaling Technologies, 1:1000), ACC1(rabbit #3676, Cell Signaling Technologies, 1:1000), or β-actin (mouse A5441, Sigma, 1:2000). Secondary antibodies were IRDye800CW goat-anti-mouse IgG (926-32210, LiCor Biosciences) and IRDye 680LT goat-anti-rabbit IgG (926-68021, Li-Cor Biosciences) both at 1:5,000. Immunoblots were imaged and analyzed with the Odyssey Infrared Imaging System and Image Studio Lite (LiCor Biosciences).

### Cell proliferation

Real-time imaging (IncuCyte, Sartorius, Ann Arbor, MI) was used to measure proliferation of cells at × 10 magnification. UCD4 cells were plated at 30,000 cells/well, T47D cells at 20,000 cells/well, sextuplicate samples in 96-well plates. For treatments, cells were given vehicle, 10 μM TVB-2640 plus 1 μM 4OH-Tam, or 100 nM Fulv on day one. Cells were counted under phase contrast immediately after treatment and subsequently taken every 4 h for 6 days. Counts were calculated as fold change over time zero. Significance was assessed by two-way repeated measures ANOVA for growth curves and t-tests for individual time points.

### Statistical Analyses

Statistical analyses were performed with GraphPad Prism 10. Data were plotted as means ± SEM. Two-tailed p <0.05 was considered statistically significant. Asterisks denote levels of statistical significance as determined by one or two-way ANOVA with Tukey’s or Dunnett’s multiple comparisons test, unpaired two-tailed t-test, one-sided Fisher’s exact test, log- rank Mantel-Cox test, Wilcoxon rank sum test, or Chi-squared test for trend. Specific tests used are indicated in the figure legends. All experiments were performed in minimum triplicate biological replicates unless noted otherwise.

## Results

### Lipid metabolic pathways are upregulated with endocrine resistance

Endocrine resistant cell lines exhibited reduced proliferation rates compared to their parental counterparts (**Figure S1a-b**). Both T47D and UCD4 endocrine resistant cell lines retain ER expression, albeit at varying levels (**Figure S1c)**. To investigate transcriptional changes associated with resistance, RNA-sequencing was performed on parental and endocrine resistant T47D (TamR, FulvR) and UCD4 (TamR, FulvR, EWD) cells. Principle component analysis revealed distinct gene expression profiles between the parental and endocrine resistant cells, as well as among the different resistant variants (**Figure 1a**). Gene Ontology (GO) analysis of the Biological Processes category identified numerous significantly up- and downregulated terms, with lipid metabolism and associated processes emerging as commonly altered features across the endocrine resistant cell lines (**Figure 1b**). Heatmaps of shared up- and downregulated genes in the Response the Lipid category in T47D (**Figure 1c**) and UCD4 (**Figure 1d**) endocrine resistant cells are depicted. Notably, lipid metabolism was consistently altered, with more pathways and genes upregulated than down regulated, highlighting the role of lipid metabolic reprogramming in endocrine resistance.

**Figure 1:**
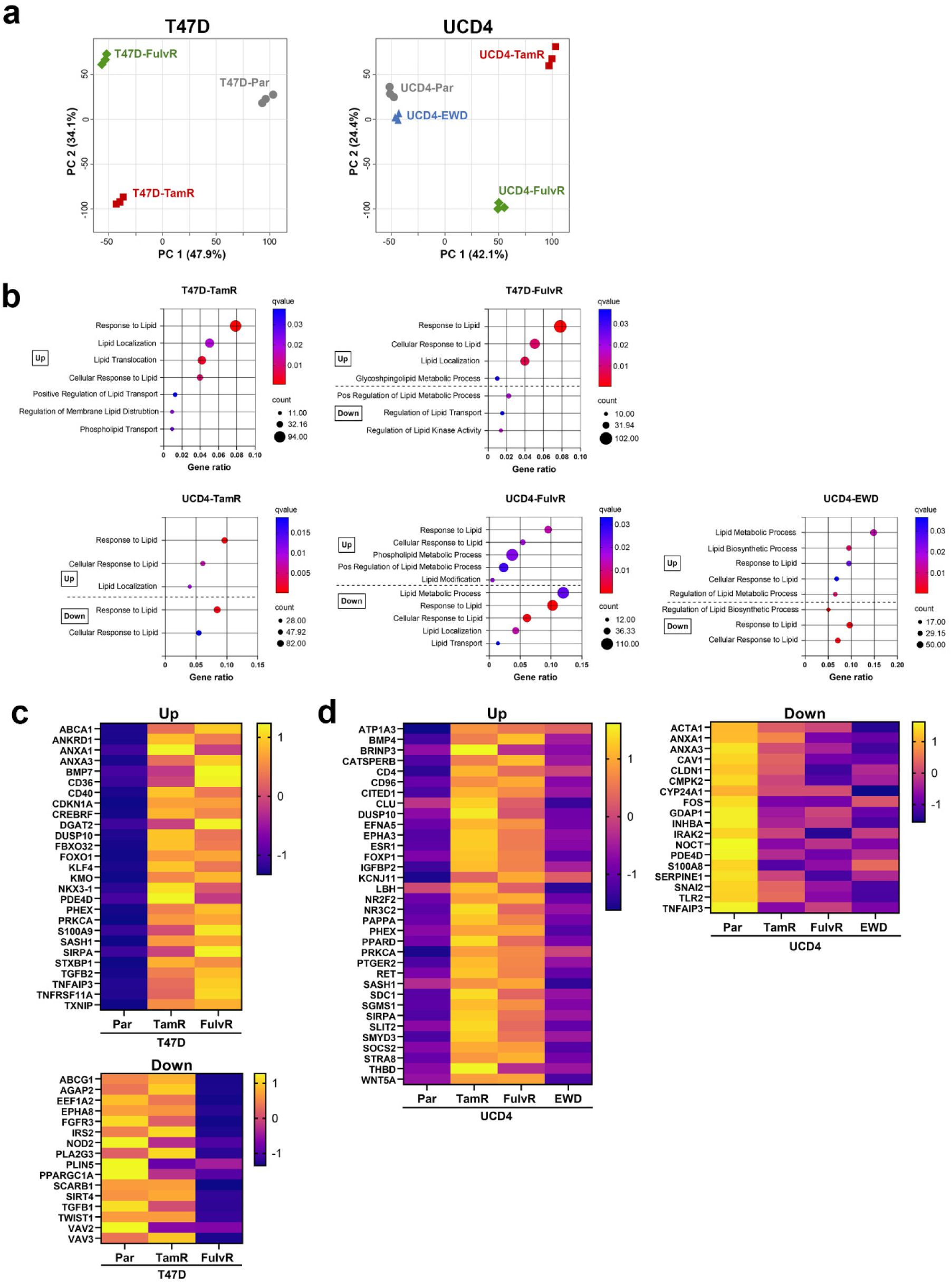
Lipid metabolic pathways are upregulated in endocrine resistant breast cancer cells. a) Principal component analysis (PCA) plots of T47D and UCD4 parental (Par) and endocrine resistant lines based on RNA-sequencing data (triplicate samples). Over representation analysis (ORA) of significantly up- and downregulated genes in lipid pathways (q<0.05, log2 fold change ≥ 2) from T47D and UCD4 endocrine resistant cells using Gene Ontology (GO) Biological Processes gene sets. T47D-TamR cells showed no significantly downregulated lipid pathways. In some cases, gene cohorts were both significantly up- and downregulated. c and d) Heatmaps of significantly up- and downregulated genes from the “Response to Lipid” process, shared between, (c) T47D-TamR and -FulvR cells (upregulated) and T47D-FulvR cells (downregulated), and (d) UCD4-EWD, -TamR, and -FulvR cells (both up- and downregulated).

### The global lipidome is altered in endocrine resistant breast cancer cells

Unbiased lipid mass spectrometry was performed to identify lipidomic changes associated with endocrine resistance in T47D and UCD4 parental lines, as well as their endocrine resistant derivatives. Normalized lipidomic data were analyzed using the publicly available metabolomics tool MetaboAnalyst. Similar to gene expression profiling, PCA analysis of lipidome data showed distinct clustering of samples by parental or endocrine resistant cell type (**Figure 2a-b**). Venn diagrams highlight commonly up- and downregulated lipid analytes in T47D (TamR, FulvR) and UCD4 (TamR, FulvR, EWD) cells (**Figure 2c-d)**. The top 25 significantly altered lipid analytes between endocrine resistant and parental cells are depicted in **Figure 2e-f**, with triglycerides and phospholipids among the most affected lipid species. These findings suggest that endocrine resistance in T47D and UCD4 cell lines is characterized by distinct lipidomic reprogramming, with specific alterations in triglycerides and phospholipids potentially playing a critical role in supporting the resistant phenotype.

**Figure 2:**
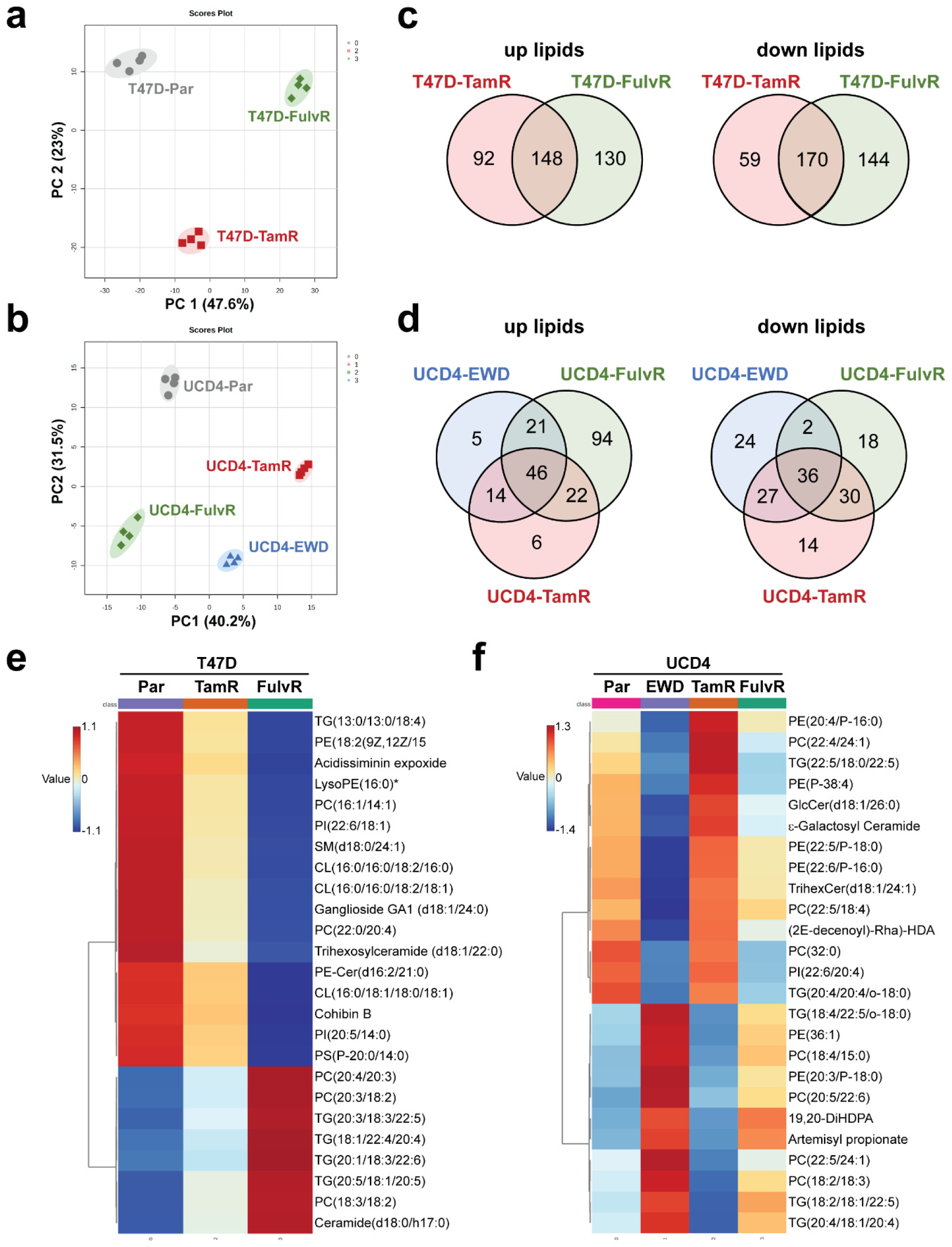
The lipid profile is altered in endocrine resistant breast cancer cells. a-b) Principal component analysis (PCA) plots of T47D (a) and UCD4 (b) parental (Par) and endocrine resistant cell samples (quadruplicate) profiled by mass spectrometry-based lipidomics. c-d) Venn diagrams of significantly altered lipid analytes (up- and downregulated, q<0.05) in T47D-TamR and -FulvR (c) and UCD4-TamR, -FulvR, and -EWD cells (d) compared to Par cells. e-f) Heatmaps of the top 25 significantly altered shared lipid analytes (up- or downregulated, PLS-DA VIP supervised clustering) between, (e) T47D-TamR and -FulvR cells, and (f) UCD4-TamR, -FulvR, and -EWD cells compared to Par cells. Plots were generated using MetaboAnalyst.

### Lipid storage is increased during endocrine resistance

To evaluate the potential function of the observed global lipidome changes, we used Lipid Ontology (LION), a publicly available resource that groups analytes within lipidomic data sets by lipid class and function (26). LION heatmap analysis on T47D and UCD4 endocrine resistant compared to parental cells clustered significant analytes into lipid signatures (**Figure S2a-b**). The lipid signature containing neutral storage lipids was the most consistently upregulated among endocrine resistant sublines in both T47D and UCD4 cell lines (**Figure 3a- b**). These include glycerolipids, triglycerides, lipid droplets, and lipid storage.

**Figure 3:**
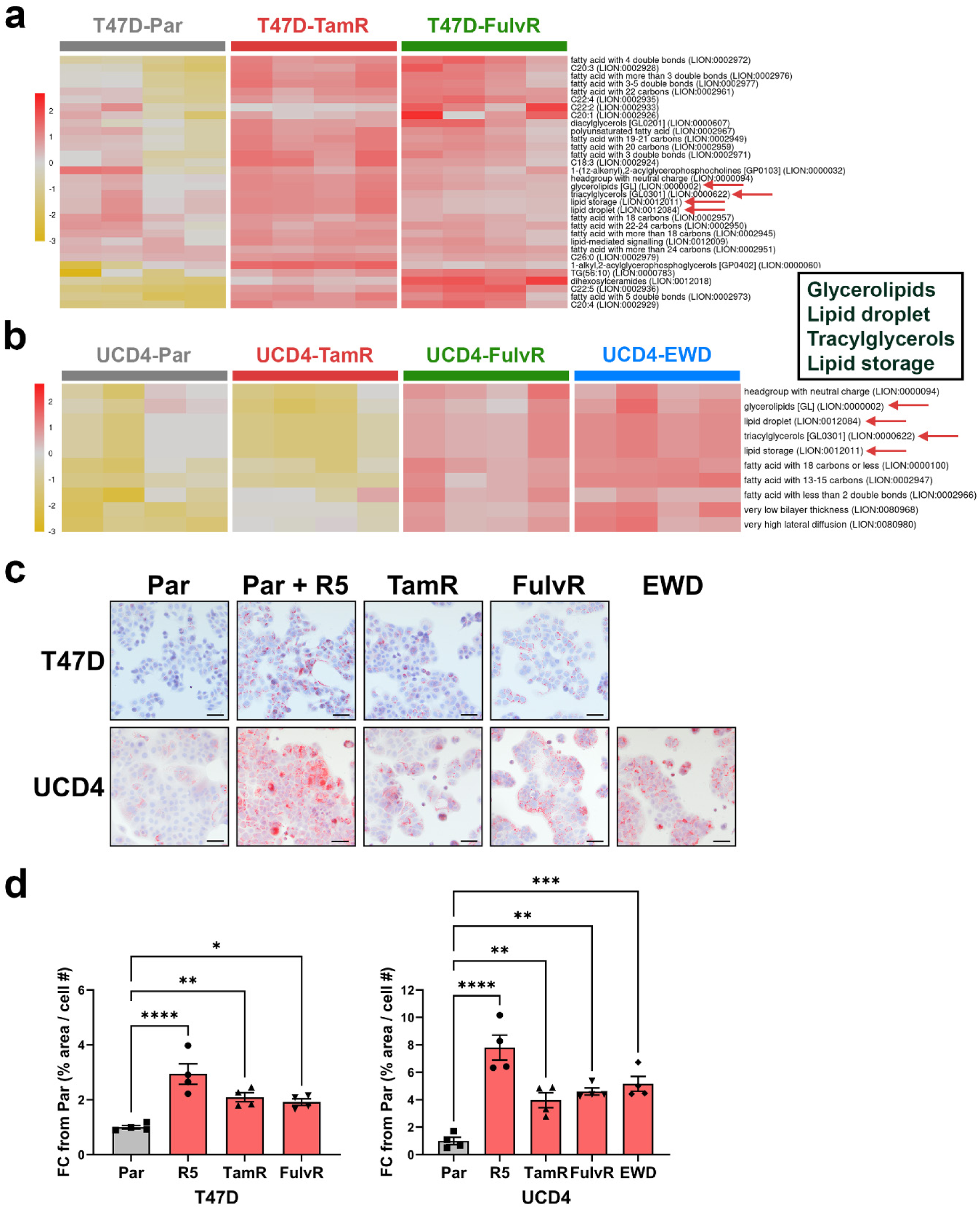
Endocrine-resistant breast cancer cells increase cytoplasmic lipid storage. a-b) Heatmaps of the Lipid Storage LION signature for T47D (a) and UCD4 (b) parental (Par) and endocrine resistant cells based on lipidomics data. Scale bars represent the mean z-score. Selected terms, with their names and IDs, are displayed on the right. Red arrows highlight analytes in the lipid storage signatures. The box on the right indicates the corresponding LION pathway annotations. c) Oil Red O (ORO) staining of T47D and UCD4 parental (Par) and endocrine resistant cells. 5 x 10^4^ cells were plated on glass coverslips in the presence of their respective drugs. Positive control cells were treated with 10 nM promegestone (R5). After 5 days, cells were formalin fixed, stained with ORO and imaged by light microscopy. Scale bars, 50 μM. d) Quantification of ORO staining using ImageJ software (4 fields per slide), expressed as ORO-positive area normalized to cell number. Fold changes from Par are shown as mean ± SEM. One-way ANOVA compares endocrine resistant to Par cells. *P<0.05; **P<0.01; ***P<0.001; ****P<0.0001.

To further explore lipid storage, we conducted Oil Red O (ORO) neutral lipid staining on parental and endocrine resistant cell lines (**Figure 3c**). A synthetic progesterone analog (promegestone, R5020) was used as a positive control for inducing LD formation (8). T47D and UCD4 parental cell lines displayed basal positive staining for LDs, which was visibly enhanced with R5020 treatment. All endocrine resistant cells showed significantly increased LD staining compared to parental controls (**Figure 3d**). Short term (72 h) treatment of T47D cells with Tam and Fulv did not significantly increase LDs, suggesting that LD accumulation is a feature of acquired resistance (**Figure S3**). To assess whether endocrine resistant cells utilize FAO, a Seahorse Palmitate Oxidation FAO assay was performed on parental and endocrine resistant cell lines (**Figure S4**). FAO was generally decreased in endocrine resistant cells, except for T47D- TamR cells, which showed a modest increase. Overall, our findings indicate that increased LD storage is a consistent feature of acquired endocrine resistance in breast cancer cells.

### TVB-2640 inhibits FASN activity and reduces lipid droplet accumulation in most endocrine-resistant breast cancer cells

Enhanced lipid storage can indicate increased FA abundance, either through the *de novo* biosynthesis or FA uptake. We assessed levels of the two rate limiting enzymes in de novo FA synthesis, FASN and Acetyl-CoA Carboxylase 1 (ACC1), by immunoblotting (**Figures 4a** **& S5** (full blot images). FASN expression was largely unchanged in endocrine resistant compared to parental cells, with an increase observed in UCD4-EWD (1.9 fold) cells. ACC1 levels were similarly unaltered in endocrine resistant cells (0.9-1.2 fold changes). Furthermore, phosphorylation of ACC1 (inactive state) did not significantly differ between parental and endocrine resistant cells (**Figure S6**), suggesting that ACC1 remains in an active state during endocrine resistance. To assess FASN activity, we performed ^13^C_2_-acetate stable isotope tracing and lipid mass spectrometry to quantify isotope incorporation into palmitate (C16), the primary product of FASN, in T47D-parental cells (with or without R5020 treatment), as well as -TamR, and -FulvR cells (**Figure 4b**). R5020-treated parental cells (positive control), -TamR, and -FulvR cells exhibited significantly higher isotope incorporation into palmitate compared to vehicle treated parental cells, indicating elevated FASN enzymatic activity endocrine resistant cells.

**Figure 4:**
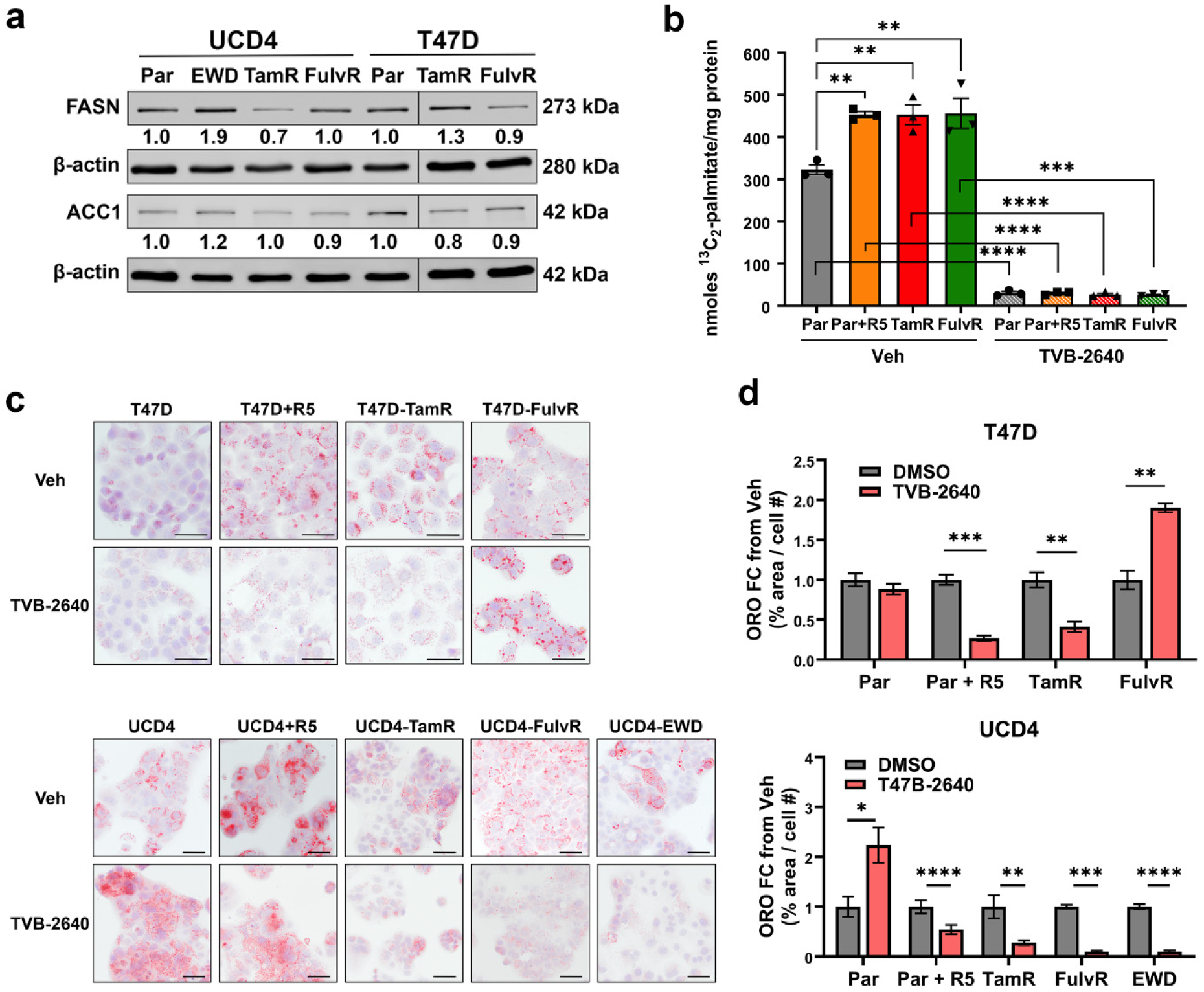
Elevated FASN activity and lipid accumulation in endocrine resistant breast cancer cells. a) Immunoblots of FASN and ACC1 in parental (Par) and endocrine resistant T47D and UCD4 cells. Bands were normalized to β-actin, and fold change over parental is indicated.

Treatment with TVB-2640 completely ablated isotope incorporation into palmitate in all tested conditions, including T47D parental (with or without R5020, -TamR, and -FulvR cells, confirming the inhibitor’s potency in blocking FASN enzymatic inhibition.

To evaluate whether FASN inhibition affects lipid accumulation, we performed ORO staining on T47D and UCD4 parental and endocrine resistant cells, with or without TVB-2640 treatment (**Figure 4c)**. In parental cells, TVB-2640 treatment blocked R5020-induced LD formation in both T47D and UCD4 cells (**Figure 4c-d**). Among the endocrine resistant derivatives, TVB-2640 significantly reduced LD accumulation in all cell lines except T47D- FulvR, where a significant increase was observed. Interestingly TVB-2640 treatment also increased LD abundance in UCD4-parental cells. Additionally, a third endocrine resistant cell line, UCD65-TamR, exhibited significantly higher LD staining compared to parental cells, which was effectively reduced by TVB-2640 (**Figure S3**). Together, these results demonstrate that while TVB-2640 is broadly effective in reducing LD accumulation, its impact is variable, depending on the cell lines and mode of resistance.

Removed lanes are denoted by a band. b) Quantification of ^13^C_2_ acetate tracer incorporation into palmitate in T47D parental (Par), -TamR, and -FulvR cells. Positive control cells were treated with 10 nM promegestone (R5) on day zero. Cells were treated with vehicle (DMSO) or 10 uM TVB-2640 for 72 h. c) ORO staining of T47D (top) and UCD4 (bottom) parental and endocrine resistant cells with or without TVB-2640. 5 x 10^4^ parental and endocrine resistant cells were plated on glass coverslips. Positive control cells were treated with 10 nM promegestone (R5) on day one. Cells were treated with vehicle (DMSO) or 10 uM TVB-2640 for 5 days. Scale bars, 50 μm. d) Quantification of ORO staining using ImageJ software (4 fields per slide), plotted as the ratio of ORO-positive area over cell number. Fold change from parental (Par) is indicated plus/minus SEM. One-way ANOVA compares endocrine resistant to parental cells. **P<0.01; ***P<0.0001; ****P<0.0001.

### Limited impact of the FASN inhibitor TVB-2640 on the growth of endocrine-resistant breast cancer cells

Since TVB-2640 effectively inhibited FASN activity, we assessed its impact on the proliferation of endocrine resistant cells. Parental and endocrine resistant derivatives of T47D and UCD4 cells were treated with TVB-2640 at the same dose used in tracing studies (10 µM) and monitored using IncuCyte live cell analysis (**Figure 5**). Minimal differences in proliferation were observed at 72 h, the timepoint used for tracing studies, with fold changes of less than 1.2 in T47D-TamR and -FulvR cells and no significant changes in UCD4 endocrine resistant compared to vehicle treated parental cells (**Figure 5a-b**). By 6 days post-treatment, proliferation of T47D cell lines was modestly reduced, with a significant >1.5 fold reduction observed in parental, TamR, and FulvR cells (**Figure 5c**). However, UCD4 cells remained largely unaffected, with no significant changes in proliferation across 6 days of treatment (**Figure 5d**). To evaluate whether higher doses of TVB-2640 could have a greater impact on cell growth, IC50 analyses were performed in T47D and UCD4 parental and endocrine resistant cells (**Figure S8a-b**). T47D cells displayed IC50 values ranging between 9-19 µM, indicating sensitivity at doses slightly above the tracing study concentration. UCD4 cell lines did not achieve efficient cytotoxicity at the highest solvent-compatible dose tested, and IC50 values could not be determined. Similarly, UCD65 cells exhibited no significant sensitivity to TVB-2640, except at the highest dose tested (100 µM) (**Figure S8c**). These findings demonstrate that while TVB-2640 effectively inhibits FASN activity and frequently reduces LD accumulation, these effects do not consistently translate into significant growth inhibition in endocrine resistant breast cancer cells.

**Figure 5:**
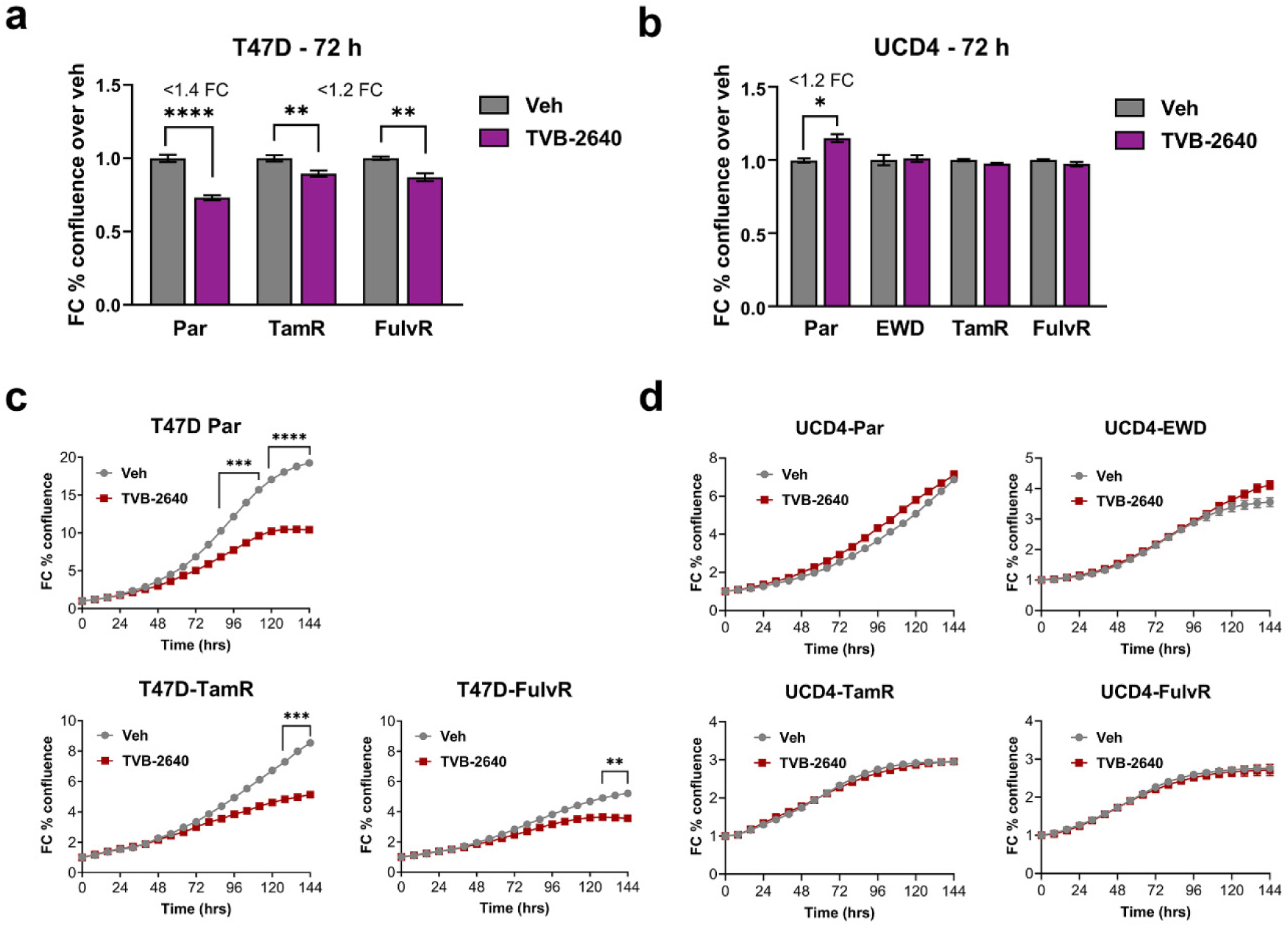
Minimal impact of the FASN inhibitor TVB-2640 on endocrine resistant breast cancer cell growth. a-b) fold change (FC) bar graphs of percent confluence at the 72 h timepoint for T47D (a) and UCD4 (b) parental (Par) and endocrine resistant cells treated with vehicle (Veh, DMSO) or 10 uM TVB-2640. FC is indicated where significant, based on a t-test comparing treatment to Veh; *P<0.05; **P<0.01; ****P<0.0001. c-d) Proliferation growth curves of T47D (c) and UCD4 (d) parental (Par) and endocrine resistant cells treated with vehicle (DMSO) or 10 uM TVB-2640 over 6 days. FC from timepoint 0 was calculated using the IncuCyte imaging system. Two-way ANOVA significance was calculated, with significance indicated at timepoints with ≥1.5 FC. **P<0.01, ***P<0.001; ****P<0.0001.

### Endocrine resistance increases lipids with unsaturated acyl chains

Our LION enrichment analysis revealed a higher abundance of lipids with increased desaturation, particularly evident in T47D endocrine resistant cells (**Figure S2**). To explore this in greater detail, significantly altered lipid analytes were categorized based on their degree of desaturation, defined as 6-12 double bonds (**Figure 6a**). The top ten up- and downregulated lipid species are for T47D (TamR, FulvR) and UCD4 (TamR, FulvR, EWD) cells compared to parental cells are shown. In general, highly desaturated lipids were more frequently upregulated than downregulated in endocrine resistant cells. To compare the absolute abundance of desaturated lipids, cumulative lipid species detected by lipidomics were stratified by the of unsaturation (≥6 double bonds) (**Figure 6b**). T47D-TamR and -FulvR cells showed significant increases in total lipid species with high degrees of unsaturation, particularly those with ≥10 double bonds, while UCD4 endocrine resistant cells exhibited a more bivalent trend. Lipid species with lower saturation were significantly decreased, whereas complex lipids with ≥11 double bonds were consistently significantly increased.

**Figure 6:**
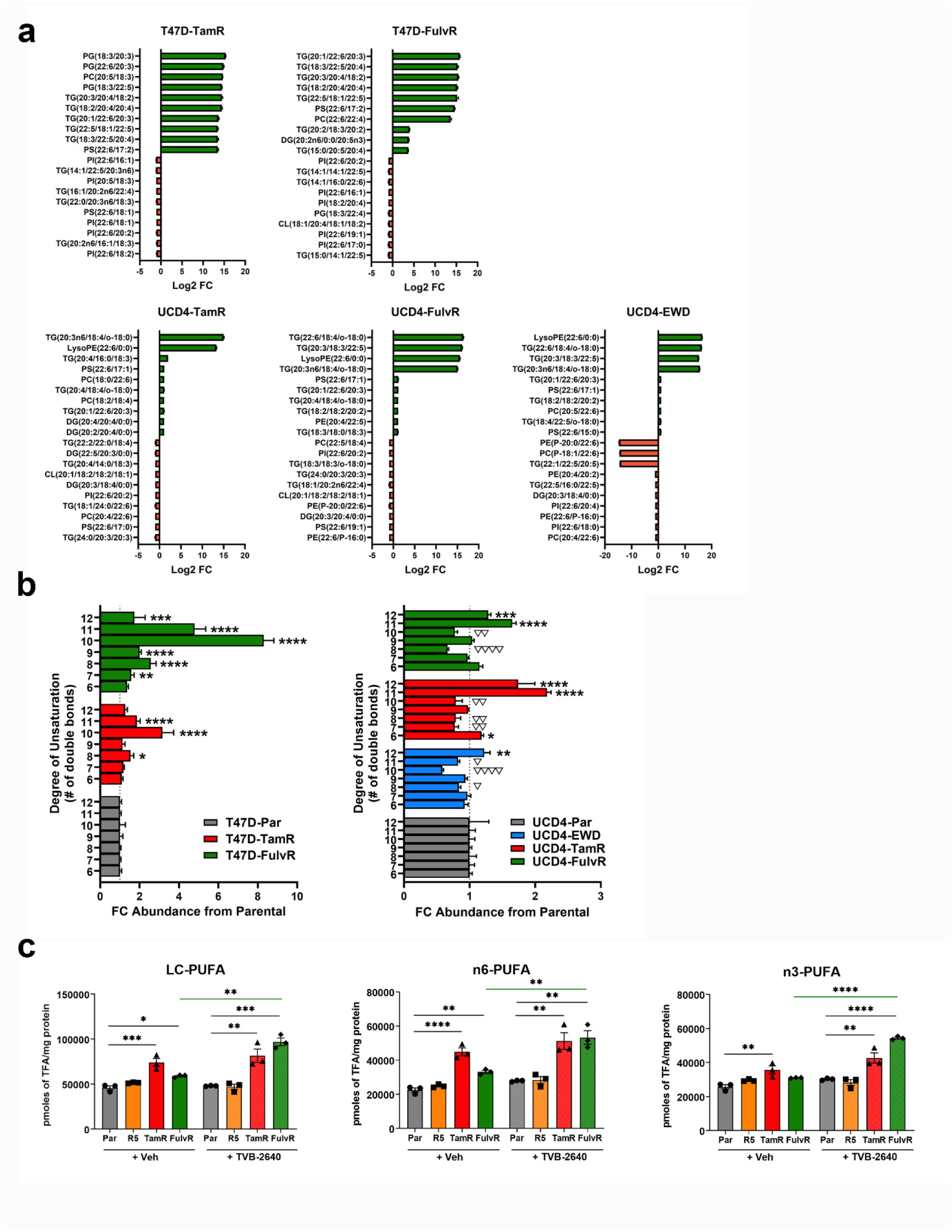
Lipid analytes with highly unsaturated acyl chains are increased in endocrine resistant breast cancer cells. a) Top ten upregulated and downregulated lipid analytes with ≥6 double bonds in T47D (top) and UCD4 (bottom) endocrine resistant cells compared to parental cells. Log2 fold changes ± SEM are shown, with species listed on the left. b) Fold changes in absolute concentrations of lipid analytes grouped by degree of saturation (≥6 double bonds) in T47D (left) and UCD4 (right) endocrine resistant cells compared to parental (Par). One-way ANOVA was applied to compare endocrine resistant to parental cells. Upregulated are indicated by *P<0.05; **P<0.01; ***P<0.0001; ****P<0.0001. Downregulated are indicated by ▽P<0.05, ▽▽P<0.01, ▽▽▽▽P<0.001. c) Concentration of long chain (LC), n-6, and n-3 PUFAs in T47D-TamR and -FulvR compared to parental cells, treated with vehicle (Veh, DMSO) or 10 uM TVB2640 for 72 h. Positive control cells were treated with 10 nM promegestone (R5) on day zero. One-way ANOVA compared T47D-TamR and -FulvR, and R- treated cells to Par T47D cells within the Veh and TVB-2640 treated groups. T-tests (green lines) indicate compare treatments ± TVB-2640. *P<0.05; **P<0.01; ***P<0.0001; ****P<0.0001.

To further investigate PUFA abundance, we analyzed PUFA levels in T47D parental, - TamR, and -FulvR cells from our tracing studies, where acyl chains were chemically liberated. Long chain (LC) PUFAs (≥18 carbons), n-6 PUFAs (e.g. arachidonic acid, AA) and n-3 PUFAs (e.g eicosapentaeonic acid, EPA, and docosahexanoic acid, DHA) were significantly increased in T47D-TamR and -FulvR compared to parental cells (**Figure 6c**). Interestingly, FASN inhibition with TVB-2640 further increased LC, n-3, and n-6 PUFA levels, specifically in T47D-FulvR cells, which also exhibited increased lipid accumulation following drug treatment (**Figure 4c-d**). No differences in LC, n-6, or n-3 PUFA levels were observed with R5020 treatment, either in the presence or absence of TVB-2640, indicating that elevated PUFA synthesis is a unique feature of endocrine resistant breast cancer cells. Together, these findings demonstrate that LC-PUFAs are specifically increased in endocrine resistant compared to parental cells, implicating desaturase activity as a potential alternate target in drug-resistant breast cancer.

## Discussion

Lipid metabolic reprogramming is a hallmark of breast cancer and varies across its major subtypes: ER+, HER2+, and TNBC (27). In this study, we focused on aberrant lipid metabolism in ER+HER2− breast cancer during acquisition of endocrine resistance. We found that endocrine resistant breast cancer cells, compared to their sensitive counterparts, commonly increase lipid storage and exhibit lipidomes enriched with PUFAs with ≥6 double bonds. Our data suggest that endocrine resistant cells enhance both de novo FA synthesis and the processing of exogenous FAs into PUFAs. These findings highlight that breast cancer cells adapt to chronic endocrine disruption by activating multiple FA synthesis and processing pathways, and demonstrate that targeting synthesis alone is unlikely to be effective in ER+ endocrine resistant breast cancer.

LD accumulation is a common feature of aggressive cancers, although their precise role in tumor progression remains unclear (28). LDs are organelles that are synthesized in the endoplasmic reticulum and are mainly composed of neutral lipid triglycerides and cholesterol esters. These LDs provide triglycerides for energy reserves, membrane synthesis, and signaling. A novel role for LDs in aggressive cancers is their ability to sequester drugs; Schlaepfer et al demonstrated that progestins increase LDs in breast cancer cells, which then sequester taxanes (8). We speculate that LDs may also sequester Tam and Fulv in endocrine resistant cells as both are fat soluble molecules. TamR breast cancer cells have previously been observed to increase LDs (12). Our studies demonstrate that resistance to Tam, Fulv, and EWD are all accompanied by exacerbated lipid storage, suggesting that increased LD production may be a common stress adaptation to chronic endocrine disruption. While TNBC cells typically have higher levels of FAO compared to ER+ cells (29), Ahn et al demonstrated that ER+ TamR breast cancer cells have a FAO gene signature resembling that of TNBC (30). Using a Seahorse FAO assay, we detected a marginal increase in FAO in T47D-TamR cells, with T47D-FulvR and UCD4 endocrine resistant lines showing decreased FAO compared to parental cells. Overall, LDs accumulate during endocrine resistance, and these structures have multifaceted roles in FA metabolism and drug resistance.

It is well recognized that breast cancer versus normal cells increase FASN driven de novo FA synthesis (6). FASN is a long-studied lipid metabolism target as it is universally increased in breast carcinoma cells and linked to worse prognosis (15, 31, 32). Most preclinical studies have found FASN blockade synergizes with chemotherapy drugs (16). However, investigation of FASN inhibition in the context of endocrine resistance is limited. TVB compounds are reversible imidazopyridine-derived inhibitors of FASN. Since endocrine resistant cells have elevated FASN levels in tandem with increased lipid storage, we tested how the new generation FASN inhibitor TVB-2640 impacted endocrine resistant cells. TVB-2640 effectively decreased FASN activity demonstrated by lack of palmitate synthesis, and mostly decreased LD abundance in treated cells. An exception was T47D-FulvR cells that increased LDs with FASN blockade. In T47D- FulvR cells we noted FASN inhibition increases LC-PUFAs. This suggests that some cells may react to blockade of de novo synthesis by processing essential imported FAs such as alpha-linolenic acid into LC-PUFAs. FASN blockade reduced the growth of all T47D cell lines by 5-6 day but not at 3-day time points, indicating a delayed effect, and one that does not coincide universally with LD reduction. TVB-2640 treatment reduced LDs in all UCD4 endocrine resistant cells and UCD65-TamR cells. Interestingly, growth of UCD4 cells was unaffected by TVB-2640, and UCD65 only at cytotoxic doses, suggesting some ER+ cells are not reliant on FASN activity and LDs for their drug resistance. LDs alone are therefore not a reliable read-out of FASN activity. Both the Phase I trial of TVB-2640 and the ongoing Phase II trial (prioritizing advanced HER2+ breast cancers) are in combination with chemotherapy (NCT03179904).

Malonyl carnitine, a metabolite of malonyl-CoA, increased in the serum of patients taking TVB- 2640, demonstrating in vivo target efficiency (17). The efficacy of TVB drugs in combination with endocrine resistance, despite effective FASN blockade, may have other off target activities. For example, Gruslova et al demonstrated that the closely related FASN inhibitor TVB-3166 decreased growth of MCF7-TamR cells potentially through degradation of ER (33). Our data insinuate that blocking de novo FA synthesis in endocrine resistant breast cancer cells may not be sufficient to improve treatment due to well recognized lipid metabolic flexibilities. A recent study found that inhibiting ACC1 to block malonyl-coA synthesis could resensitize long term estrogen deprived breast cancer cells in some contexts, prospectively by blocking peroxisome mediated LC-FA oxidation (34).

Our experiments uncovered that endocrine resistant versus parental cells contain a higher content of lipids with increased degrees of unsaturation, particularly PUFAs with greater than nine double bonds. Interestingly, in T47D-FulvR cells, PUFAs increased further upon FASN inhibition. PUFAs provide building blocks for membrane expansion, their synthesis provides energy in the form of NADPH for other metabolic reactions, and they serve as substrates for the synthesis of oxidized signaling lipids. However, excessive storage of PUFAs leaves cells vulnerable to lipid peroxidation, which can trigger cell death through processes such as ferroptosis (35). The enzymes that catalyze the generation of double carbon bonds in FAs are the FA desaturases (FADS). The main FADS in humans include FADS5, otherwise known as Stearoyl-CoA Desaturase (SCD), which produces monounsaturated FAs from C16 and C18 FAs, and FADS1, FADS2, that produce n-6 and n-3 PUFAs from the essential C18 FAs linolenic acid and α-linolenic acid, respectively (36). While most studies have focused on inhibiting SCD, several recent studies have focused on blocking other FADS in cancer. Vriens et al identified that by inhibiting SCD in breast cancer cells (not endocrine resistant), FADS2 compensates by producing the alternative FA sapienate, and FADS2 inhibition could reverse this process (31).

Thus, disrupting PUFA synthesis by FADS could be particularly damaging to endocrine resistant cells. We observed other lipid metabolic changes in endocrine resistant cells such as altered membrane lipid content that could also reveal vulnerabilities in endocrine resistance and will require further investigation

## Conclusions

We demonstrate that breast cancer cells resistant to endocrine therapies have increased FASN activity, PUFAs with >6 double bonds, and lipid storage. Targeting de novo synthesis in the presence of endocrine drugs selectively impacted cell growth and did not correlate with LD reduction, suggesting endocrine resistant breast cancer cells may evade FASN inhibition through other lipid metabolic pathways. This prospectively includes the uptake of essential PUFAs that undergo further elongation and desaturation events and may present a unique vulnerability for targeting desaturation in endocrine resistant breast cancer cells.

## Abbreviations

ACC: Acetyl CoA carboxylase
ER: Estrogen Receptor alpha
EWD: Estrogen withdrawal
FA: Fatty acid
FADS2: Fatty Acid Desaturase 2
FASN: Fatty acid synthase
FAO: Fatty acid oxidation
Fulv: Fulvestrant
FulvR: Fulvestrant resistant
PUFA: Polyunsaturated fatty acid
LC-PUFA: Long chain polyunsaturated fatty acid
LD: Lipid droplet
ORO: Oil Red O
PR: Progesterone receptor
SERD: Selective estrogen receptor degrader
SERM: Selective estrogen receptor modulator
SREBP1: sterol regulatory element-binding transcription factor 1
Tam: Tamoxifen
TamR: Tamoxifen resistant
TNBC: Triple negative breast cancer

## Declarations

### Ethics approval and consent to participate

All animal procedures were approved by the Colorado Institutional Animal Care and Use Committee.

## Acknowledgements

We thank the University of Colorado School of Pharmacy Mass Spectrometry Facility for their consultation and services. We thank the University of Colorado Cancer Center Bioinformatics and Biostatistics Shared Resource supported by P30CA046934.

## Author contributions

A.V.W, P.K., M.C.R., and C.A.S. designed the studies. R.R.V., K.B.H., and M.C.R. conducted lipidomic and tracing studies. A.V.W., D.R., and K.E.C. conducted cell culture experiments. A.V.W, D.R., K.E.C., A.E.L., R.R.V., M.C.R. and C.A.S. conducted all data analysis. A.V.W. and

C.A.S. drafted manuscript. D.R., J.F.S, H.M.B., R.R.V., P.K., and M.C.R. reviewed and edited manuscript. The project was supervised and funded by M.C.R. and C.A.S. All authors have read and agreed to the published version of the manuscript.

## Availability of data and materials

The raw RNA-sequencing data is available in the NCBI Gene Expression Omnibus (GEO Accession GSE281935). The unique cell lines are available to qualified individuals for research purposes through formal request, MTA agreement, and appropriate fee.

## Consent for publication

Not applicable.

