## Supplemental Figures for "Lipid metabolic reprogramming drives triglyceride storage and variable sensitivity to FASN inhibition in endocrine-resistant breast cancer cells"

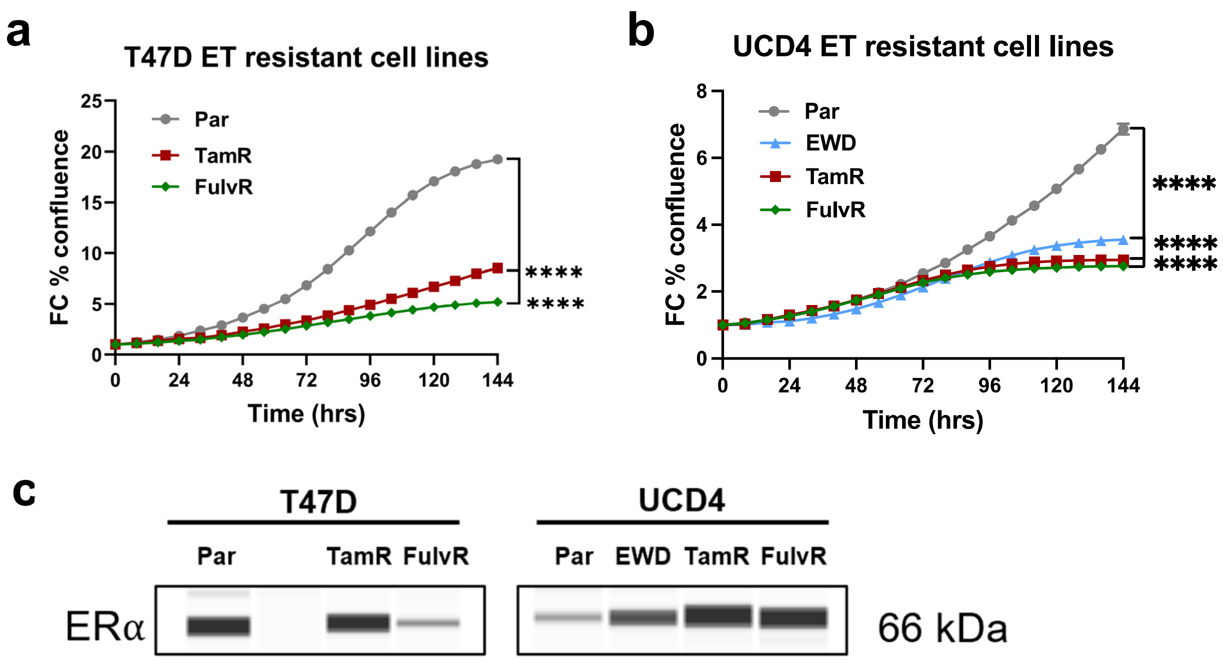


**Supplemental Figure S1: Growth and ER expression in endocrine resistant cell lines.** a-b) Proliferation growth curves of parental and endocrine resistant T47D (a) and UCD4 (b) cells. Fold change (FC) from timepoint 0 was calculated using the IncuCyte platform. T47D and UCD4 cell lines were plated at 7500 and 15,000 cells/well, respectively, in sextuplicate, and images were captured every 4 h for 6 days. Statistical significance at the final timepoint was calculated using One-way ANOVA. ****$P<0.0001$. c) JESS ProteinSimple analysis of ER (alpha) expression in T47D and UCD4 cells. Bands represent peak intensities normalized to total protein, as analyzed using JESS software.

a

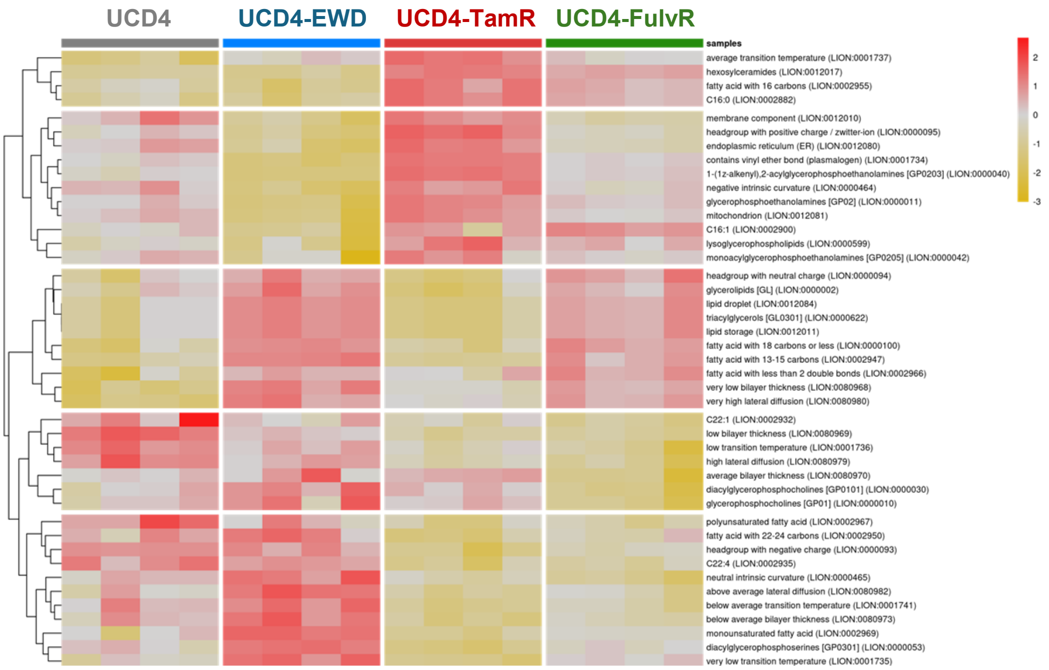


b

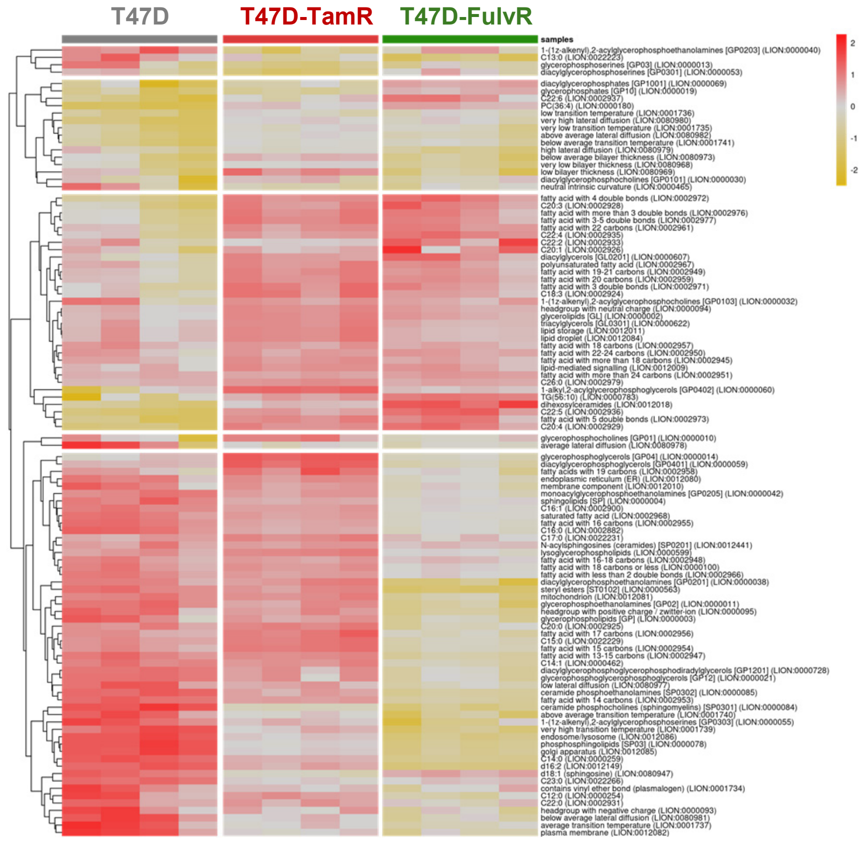


**Supplemental Figure S2: Full LION heatmaps of endocrine resistant breast cancer cells.** Full LION heatmaps and hierarchical clustering of lipidomics data from T47D (a) and UCD4 (b) parental and endocrine resistant cell lines. Lipid signatures are grouped into five clusters based on hierarchical clustering. Heatmap colors (yellow to red) represent mean z-scores. Names and IDs of the selected lipid terms are displayed on the right.

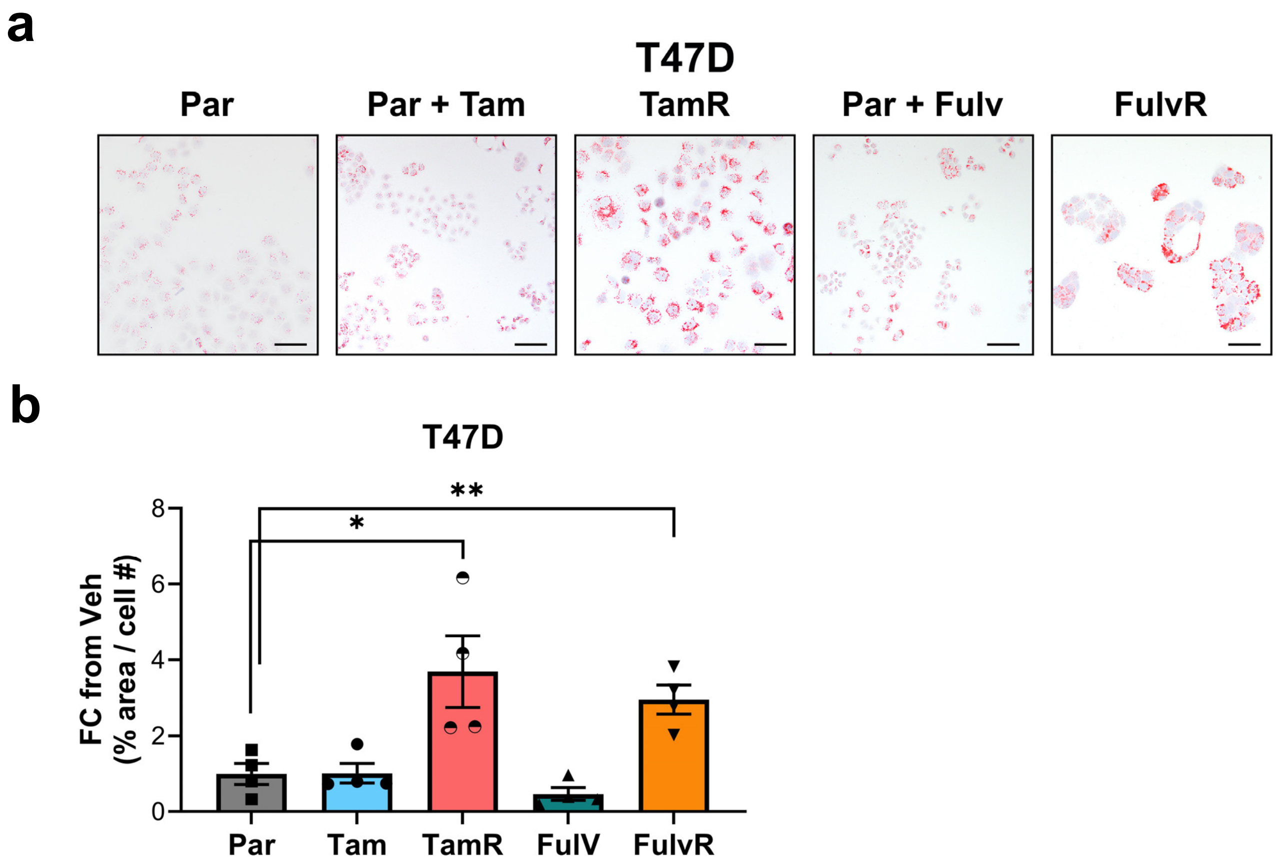


**Supplemental Figure S3. Lipid droplet accumulation in T47D cells with acute endocrine drug treatment**. a) Oil Red O (ORO) staining of T47D-parental cells treated with vehicle (Par), 1 μM 4-OH-Tam, or 100 nM Fulv for 72 h, compared to TamR and FulvR cells. A total of 5 x $10^4$ cells per condition were plated on glass coverslips with the appropriate drug treatments. Cells were formalin fixed, stained with ORO and imaged by light microscopy. Scale bars, 50 μM. b) Quantification of ORO staining. Images were analyzed using ImageJ software (4 fields per condition) to calculate the ORO-positive area normalized to cell number. Fold change relative to T47D-Par is shown. Statistical comparisons were performed using one-way ANOVA to compare endocrine resistant to parental cells. *P<0.05; **P<0.01.

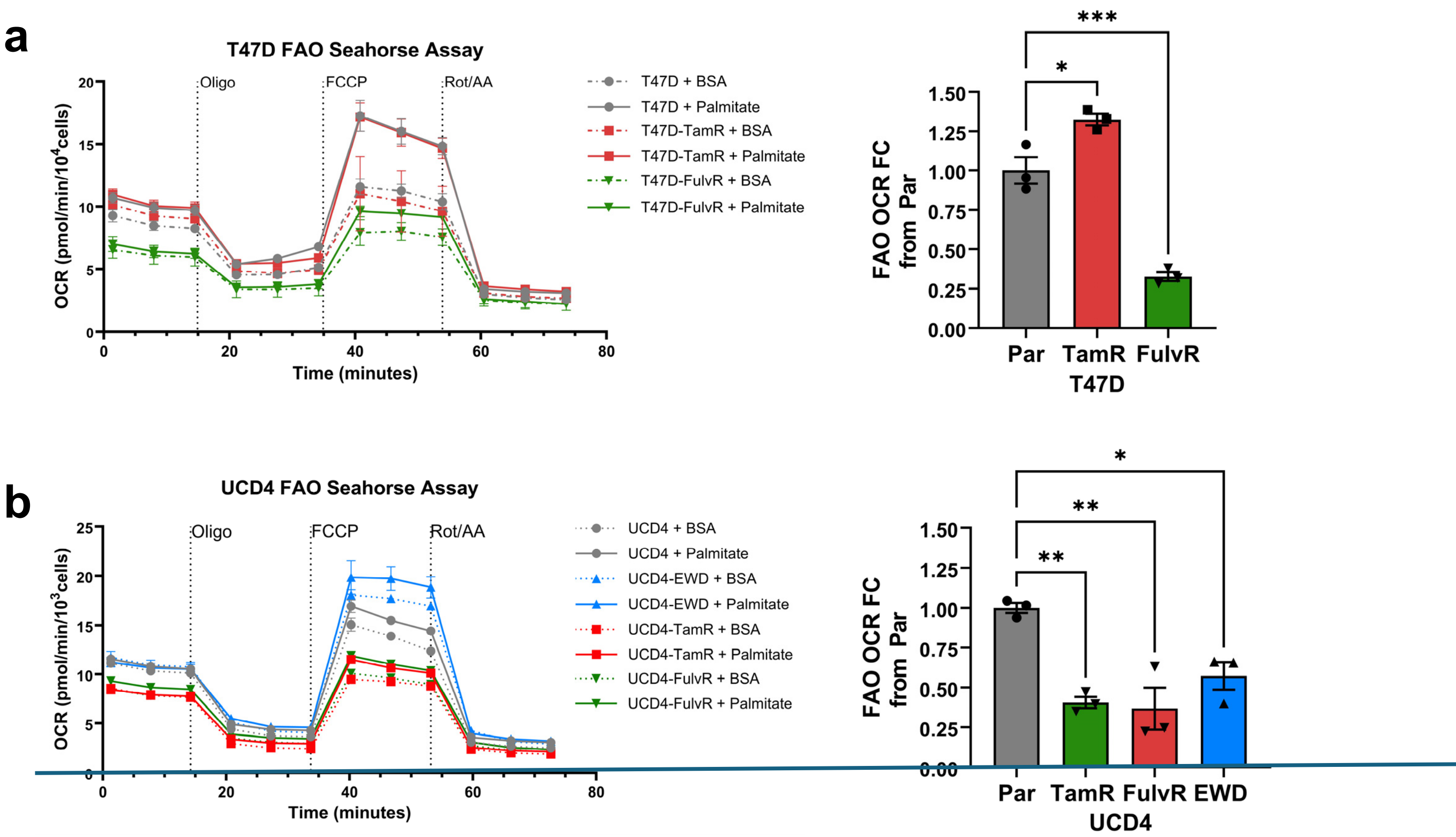


**Supplemental Figure S4. Fatty acid oxidation (FAO) in endocrine resistant cells.** a-b) oxygen consumption rate (OCR) kinetic curves. 7,500 T47D (a) and 15,000 UCD4 (b) parental and endocrine resistant cells were plated in XF96-well Seahorse Assay plates (n=6 each), nutrient deprived, and supplemented with BSA control or XF Palmitate-BSA conjugate 4 hours prior to performing the assay. Assays were standard XF Mito Stress Tests conducted in a 96XFe format. OCR was normalized to cell count obtained by Hoechst 33342 dye and Agilent Cytation instrument post-assay. Oligo = Oligomycin; FCCP = Trifluoromethoxy carbonylcyanide phenylhydrazone; Rot/AA = Rotenone and Antimycin A. Error bars represent SEM. c-d) Relative FAO-specific OCR rates in T47D (c) and UCD4 (d) parental (Par) and endocrine resistant cells. FAO-OCR was determined by comparing palmitate versus BSA conditions minus non-mitochondrial respiration. One-way ANOVA compared to Par is shown. *P<0.05, **P<0.01, ***P<0.001,

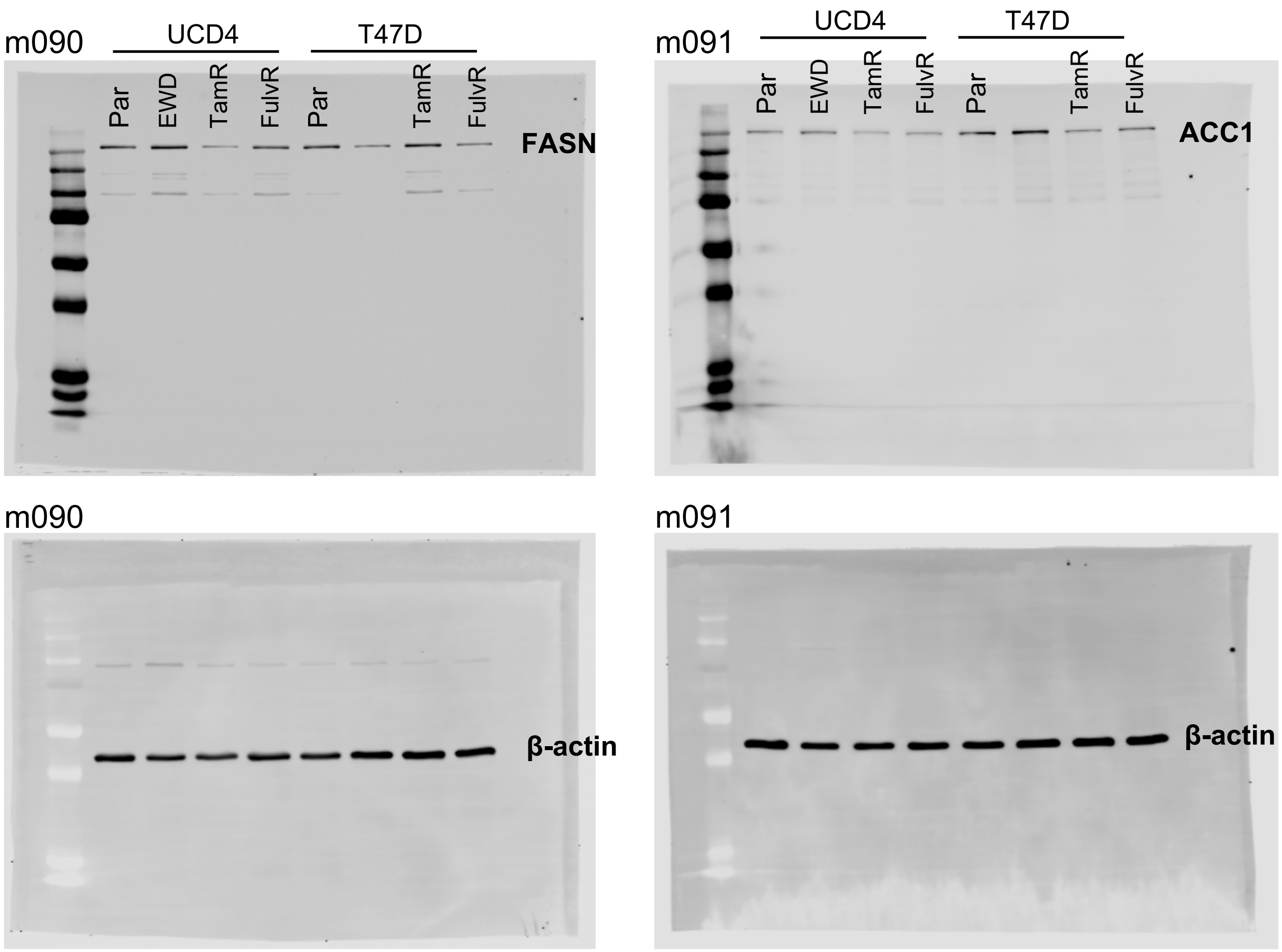


**Supplemental Figure S5. Full immunoblots of FASN and ACC1 in endocrine resistant breast cancer cells.** Full immunoblots for FASN and ACC1 protein levels in parental (Par) and endocrine resistant T47D and UCD4 cells, corresponding to the cropped blots shown in Figure 4a. β-actin used as loading control (bottom). Note that some lanes were not included in this study.

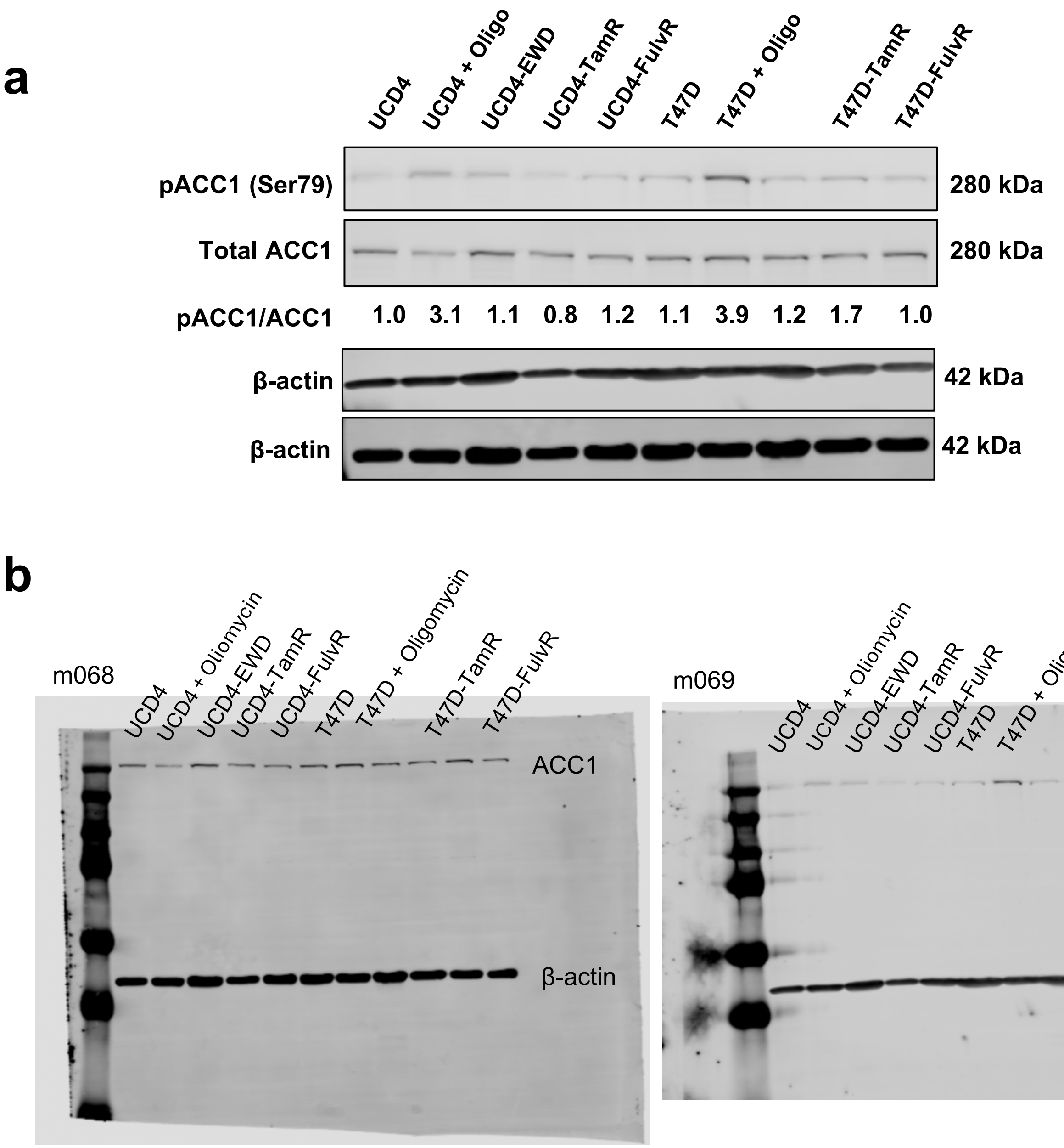


**Supplemental Figure S6. Endocrine resistant cell lines retain activated ACC1.** a) Immunoblot analysis of phosphorylated ACC1 (pACC1 at Ser79) and total ACC1 levels in parental and endocrine resistant T47D and UCD4 cells. β-actin was used as a loading control. Protein bands were normalized to β-actin, and the fold change of pAAC1/ACC1 relative to parental cells is indicated. b) Full immunoblots corresponding to data depicted in (a). Note that some lanes were not included in this study.

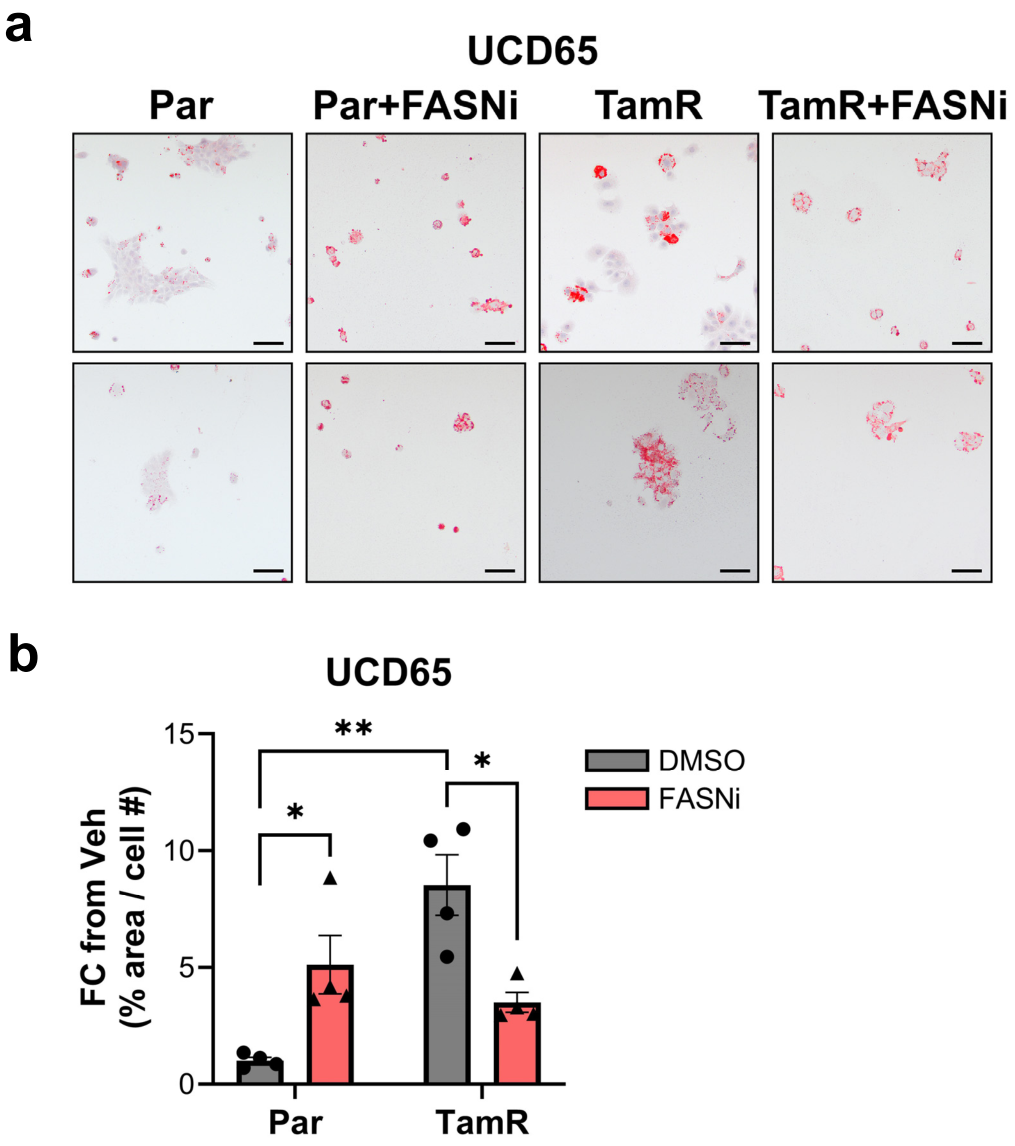


**Supplemental Figure S7. Lipid droplet accumulation in UCD65 parental and TamR cells treated with TVB-2640.** a) Oil Red O (ORO) staining of UCD65-parental (Par) and TamR cells. A total of 7.5 x $10^4$ cells per condition were plated on glass coverslips in standard media. Cells were treated with vehicle (DMSO) or 10 uM TVB-2640 for 72 h. Cells were formalin fixed, stained with ORO, and imaged by light microscopy. Scale bars, 50 μM. b) Quantification of ORO staining. Images were analyzed by ImageJ software (4 fields per condition) to determine the ORO-positive area normalized to cell number. Fold change relative to Par cells is shown. Statistical comparisons were performed using one-way ANOVA. *P<0.05; **P<0.01.

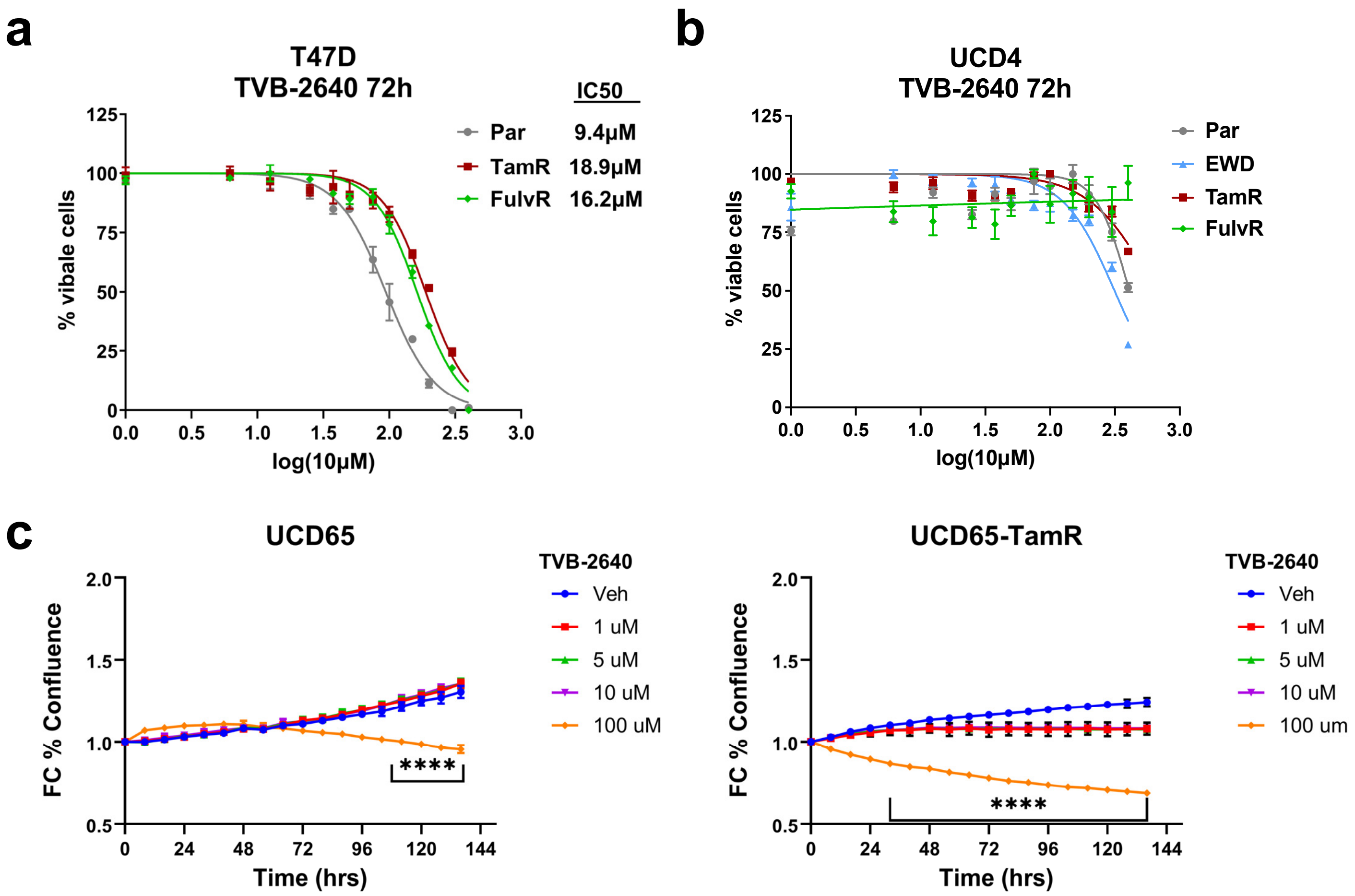


**Supplemental Figure S8. Dose curves for TVB-2640 for endocrine therapy-resistant breast cancer cells.** a-b) Dose-response curves of T47D (a) and UCD4 (b) parental and endocrine therapy-resistant cells treated with increasing concentrations of TVB-2640 or vehicle (DMSO). The IC50 value for TVB-2640 is indicated on the graph for T47D cells. The IC50 for UCD4 cells could not be determined, as the drug was insufficient to induce significant cell death even at the highest concentration tested. c) Growth curves of UCD65-Par and -TamR cells treated with increasing concentrations of TVB-2640. Fold change (FC) in percent confluence relative to parental cells is shown. Data represent sextuplicate samples ± SD. Statistical significance for differences in fold changes greater than 1.2 was calculated using a two-way ANOVA; significance is indicated. ****P<0.0001.
