## Supplemental Methods for "Lipid metabolic reprogramming drives triglyceride storage and variable sensitivity to FASN inhibition in endocrine-resistant breast cancer cells"

**Supplementary Materials**

**Supplemental Methods**

**Growth Analysis**

Real-time imaging (IncuCyte, Sartorius, Ann Arbor, MI) was used to measure proliferation of parental and ET resistant T47D and UCD4 cells at × 10 magnification. T47D cells were plated at 5000 cells per/well and UCD4 cells were plated at 15,000 cells/well in 96-well plates. Appropriate wells received maintenance concentrations of 4OHT (TamR), Fulv (FulvR), or phenol-free medium with charcoal stripped serum (EWD). Well confluence was quantified every 4 hours. Percent confluence was calculated as fold change from hour 0. Significance was assessed by one-way ANOVA/Tukey at the final time point.

**IC50 analysis**

Real-time imaging (IncuCyte) was used to measure proliferation of cells at 10x magnification. One day prior to treatment, T47D (5,000 cells/well), UCD4 (15,000 cells/well) and UCD65 (30,000 cells/well) in quintuplicate-sextuplicate in 96-well plates. The following day, serially diluted FASN inhibitor (TVB-2640) or DMSO control was added to each respective treatment group. Over the span of six days, cell proliferation in response to treatment was measured and plotted as fold change relative to time = 0 h. 72 h after treatment, growth rates stratified in a dose dependent manner. In GraphPad Prism 10, fold change values for each replicate where then plotted against transformed, log10, concentration TVB-2640 (uM x 10). Fold change values were then normalized determining 100% viability as the largest mean of each data set and 0% viability (or 100% inhibition) was determined as FC equal to or less than 1. The highest concentration of TVB-2640 (50 -100 uM) was not sufficient to induce 100% inhibition in all cell lines. Non-linear regression analysis was calculated as log (inhibitor) vs. normalized response- variable slope. All cell-lines have an R squared of 0.95 or greater.

**Protein expression analysis**

Protein levels of ER were assessed using the Jess capillary electrophoresis system (ProteinSimple, Minneapolis, MN). Protein lysates were harvested in RIPA buffer in the presence of protease and phosphatase inhibitor. Samples were diluted down to 1 mg/mL in 5x fluorescent master mix containing DTT. Samples were then denatured for 5 minutes at 95C and 0.5 μg/well loaded onto a 12-230 kDa Jess plate (SM-FL004). Lysate capillaries were subsequently probed for Estrogen Receptor alpha (rabbit, #8644, Cell Signaling Technologies), followed by a secondary chemiluminescent rabbit secondary (#DM-001). All samples were normalized to total protein using Protein Normalization reagent (DM-PN02).

**FAO Seahorse Mito Stress Assays**

For Mito Stress Test (Agilent, #103015-100) parental and ET resistant T47D or UCD4 cells were seeded into XFe96-well cell culture plates at 7.5x10^4^ cells/well and 15x10^4^ cells/well, respectively, in sextuplicate. Cells were seeded in normal assay medium and left at 1 h at room temperature (25C) before returning to the incubator (37C) to prevent edge effects. Appropriate wells received maintenance concentrations of 4OHT (TamR), Fulv (FulvR), or phenol-free medium with charcoal stripped serum (EWD). One day before the assay, a XF Sensor cartridge was hydrated with 100 µL H_2_O/well and placed in non-CO_2_ incubator (37C). 30 mL XF Calibrant was also preheated in non-CO_2_ incubator (37C). On the day of the assay, cells were nutrient deprived, then supplemented with BSA control or XF Palmitate-BSA conjugate for 4 h prior to performing the assay. Assay medium was prepared according to Agilent Assay protocol. Cell culture wells were washed twice with 100 µL assay medium and left with a final volume of 180 µL. The cell culture plate was put to de-gas in non-CO_2_ incubator (37C) for 1 h before start of assay. Oxygen consumption rate (OCR) was normalized to cell count obtained by Hoechst 33342 dye and Agilent Cytation instrument post-assay. Oligo=Oligimycin; FCCP= Trifluoromethoxy carbonylcyanide phenylhydrazone; Rot/AA=Rotenone and Antimycin A. Error bars represent SD.
